# Functional categories in macaque frontal eye field

**DOI:** 10.1101/212589

**Authors:** Kaleb A. Lowe, Jeffrey D. Schall

## Abstract

Frontal eye field (FEF) in macaque monkeys contributes to visual attention, visual-motor transformations and production of eye movements. Traditionally, neurons in FEF have been classified by the magnitude of increased discharge rates following visual stimulus presentation, during a waiting period, and associated with eye movement production. However, considerable heterogeneity remains within the traditional visual, visuomovement and movement categories. Cluster analysis is a data-driven method of identifying self-segregating groups within a dataset. Because many cluster analysis techniques exist and cluster outcomes vary with analysis assumptions, consensus clustering aggregates over multiple cluster analyses, identifying robust groups. To describe more comprehensively the neuronal composition of FEF, we applied a consensus clustering technique for unsupervised categorization of patterns of spike rate modulation measured during a memory-guided saccade task. We report ten functional categories, expanding on the traditional three. Categories were distinguished by latency, magnitude, and sign of visual response, presence of sustained activity, and dynamics, magnitude and sign of saccade-related modulation. Consensus clustering can include other metrics and can be applied to datasets from other brain regions to provides better information guiding microcircuit models of cortical function.

**SIGNIFICANCE STATEMENT:** The contribution of a brain region cannot be understood without knowing the diversity, arrangement, and circuitry of constituent neurons. Traditional descriptions of frontal eye field include visual, visuo-saccadic, and saccadic categories. Here, we employ a novel consensus clustering method to identify more reliably functional categories in neural data. While confirming the traditional categories, consensus clustering distinguishes additional, previously unappreciated diversity in neural activity patterns. Such information is necessary to formulate correct microcircuit models of cortical function.

## INTRODUCTION

Like all cortical areas, frontal eye field (FEF) is comprised of neurons distinguished by morphology, neurochemistry, biophysics, layer, and connectivity. Biophysical distinctions can be made via action potential waveforms (McCormick et al., 1985; Mitchell et al., 2007; Cohen et al., 2009; Ding and Gold, 2012; Thiele et al., 2016), calcium binding proteins (Pouget et al., 2009), and neuromodulatory receptors (Noudoost and Moore, 2011; Soltani et al., 2013). Neurons with distinct biophysical characteristics must play different roles in the cortical microcircuit (Lewis and Lund, 1993; DeFelipe 1997; Pouget et al., 2009; Zaitsev et al., 2012). Connectivity studies find FEF connected with at least 80 cortical areas (e.g., Huerta et al., 1986, 1987; Schall et al., 1993; Stanton et al., 1993; Schall et al., 1995a; Stanton et al., 1995; Markov et al., 2014), and most pyramidal neurons do not project to more than one cortical area (Markov et al., 2014; Ninomiya et al., 2012; Pouget et al., 2009). Numerous functional distinctions among FEF neurons have been reported, beginning with the traditional sorting into visual, visuomovement, and movement plus fixation and postsaccadic categories (Bruce and Goldberg, 1985; Schall, 1991). Subsequently, FEF neurons have been implicated in complex functions including visual search (Schall et al., 1995b; Thompson et al., 1996; Lee and Keller, 2008; Zhou and Desimone, 2011; Purcell et al., 2012), saccade preparation and inhibition (Hanes et al., 1998; Boucher et al., 2007; Ray et al., 2009), perceptual choice (Ding and Gold, 2012), feature-based attention (Bichot et al., 1996; Bichot and Schall, 2002; Gregoriou et al., 2009; Zhou and Desimone, 2011; Noudoost et al., 2014) transsaccadic stability (Crapse and Sommer, 2008, 2012; Shin and Sommer, 2012), planning saccade sequences (Phillips and Segraves, 2010), and anticipating reward (Glaser et al., 2016). How can so many functions be accomplished by so few neuron categories?

The problem of classification is neither new to science nor unique to neurophysiology. Cluster analysis is a powerful statistical tool, developed to find self-segregating categories in gene expression (Sharp et al., 1986), psychiatric diagnostics (Lochner et al., 2005), linguistics (Gries and Stefanowitsch, 2010), and Scotch whisky (Lapointe and Legendre, 1994). It has also been used to describe the biophysical diversity of cortical neurons (Nowak et al. 2003; Druckmann et al. 2013; Ardid et al., 2015), expanding the canonical *in vivo* description of putative excitatory and inhibitory cells. Cluster analysis should be similarly powerful for assessing the functional diversity that must parallel anatomical diversity and should reproduce the functional categories known to exist in FEF.

Cluster analysis requires strategic decisions about the method of grouping observations and how to calculate pair-wise distance, which lacks rigorous specification for clustering functional characteristics of neurons. Therefore, we applied multiple pre-processing pipelines to a large sample of FEF neurons then applied an agglomerative clustering algorithm to discover functional categories.

Because *a priori* endorsement of any particular pre-processing pipeline is impossible, and each result is unique, the results of an individual clustering procedure are difficult to interpret. However, second-order clustering procedures known as consensus clustering can reconcile outcomes from different pipelines (Strehl and Ghosh, 2002). Distinct consensus clustering procedures use different theoretical motivations and computational efficiencies (Goder and Filkov, 2008). We applied a procedure that operates on the median pairwise distance across all pre-processing pipelines because it is tractable and efficient. This consensus clustering procedure identified ten robust functional categories of FEF neurons, which elaborate functional variation among the three traditional categories plus fixation and post-saccadic neurons.

## METHODS

### Subjects and Behavioral Task

Three macaque monkeys (*M. radiata*) participated in this study. All procedures were in accordance with the National Institutes of Health Guide for the Care and Use of Laboratory Animals and approved by the Vanderbilt Institutional Animal Care and Use Committee. Monkeys were trained to perform a memory guided saccade task (Bruce and Goldberg, 1985). Trials began when a central fixation point appeared. After fixating this point for 500 ms, a peripheral target was briefly presented at 8° eccentricity at one of 8 locations separated by 45°. After a variable delay between 300 and 800 ms the fixation point was extinguished, and the monkey earned fluid reinforcement for shifting gaze to the location of the peripheral stimulus. The peripheral stimulus was re-illuminated after the saccade to provide a fixation stimulus. If the monkey maintained this fixation for 500 ms, fluid reward was delivered. If the monkey broke fixation, a 5,000 ms time-out delay occurred.

### Recording Techniques

MRI compatible headposts and recording chambers were placed over the arcuate sulcus.

Surgery was conducted under aseptic conditions with animals under isoflurane anesthesia. Antibiotics and analgesics were administered postoperatively. Details have been described previously (Schall et al., 1995a; Sato et al., 2001; Cohen et al., 2009). Electrophysiological data was obtained from linear electrode arrays, either a 24-channel Plexon Uprobe (monkeys Ga, He) or a 32-channel Neuronexus Vector Array (monkey Da). Both probes had a 150 µm recording contact spacing. Data were streamed to a data acquisition system: MNAP (40 kHz, Plexon, Dallas, TX - monkeys Da, Ga, and He) or TDT System 3 (25 kHz, Tucker Davis Technologies – monkey Da). Single units were identified online using a window discriminator (Plexon) or principle component analysis (TDT). Units recorded from the TDT system were sorted offline using Kilosort (Pachitariu et al., 2016). Eye position was collected using EyeLink 1000 (SR Research). Eye position was calibrated daily and streamed to the data acquisition system and stored at 1 kHz.

### Neuron Classification

Spike density functions (SDF) were calculated by convolving the spike trains with a function that resembles the postsynaptic influence of each spike (Thompson et al., 1996). Spike density functions were calculated only for correct trials on which the visual stimulus was presented in the visual receptive field and the saccade was made into the movement field. A sequence of classification procedures was employed. The first was based on the traditional criteria of Bruce & Goldberg (1985). A unit was considered to have visual activity if the firing rate between 50 and 150 ms after stimulus presentation was elevated more than six standard deviations above the baseline mean. A unit was considered to have movement activity if the firing rate in the 100 ms preceding the saccade was more than six standard deviations above the baseline mean and the SDF showed a positive correlation over time in the 20 ms preceding saccade. This prevents elevated delay activity with no pre-saccadic ramping from being considered movement-related activity. Visual units had visual responses with no movement activity, movement units had movement activity with no visual responses, and visuomovement units had both. Other units were considered uncategorized; we did not test for fixation or postsaccadic activity in this categorization.

Units were categorized via agglomerative hierarchical cluster analyses (Sokal and Michener, 1958; Everitt et al., 2011). These analyses iteratively combine units, or groups of units, based on the weighted average similarity of units. In each case the analysis algorithm was identical, though the method for determining similarity differed due to the scaling of discharge rates across units, method of summarizing the units’ response, or the similarity metric (Table 1; Fig. 1). The agglomerative cluster analysis was performed as follows: First, the sample was considered as *n* groups, each with one member. Then, the two groups with the smallest pairwise distance were combined into one group, leaving (*n*-1) groups, one of which with two members. The distances of this group to the other groups were determined by the weighted mean of the distances of the individuals in each group:

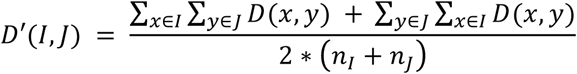

where *I* is the first group in consideration and *J* is the second, *x* are the members of group *I*, *y* are the members of group *J*, and *n*_*I*_ and *n*_*J*_ are the number of members in groups *I* and *J*, respectively. The value of 2 in the denominator is required because the distances are symmetrical and thus represented twice in the numerator of the equation. More simply, this averages the pairwise distances of the members of *I* and *J* such that the similarity of two groups will not be skewed by uneven group sizes.

**TABLE 1:**
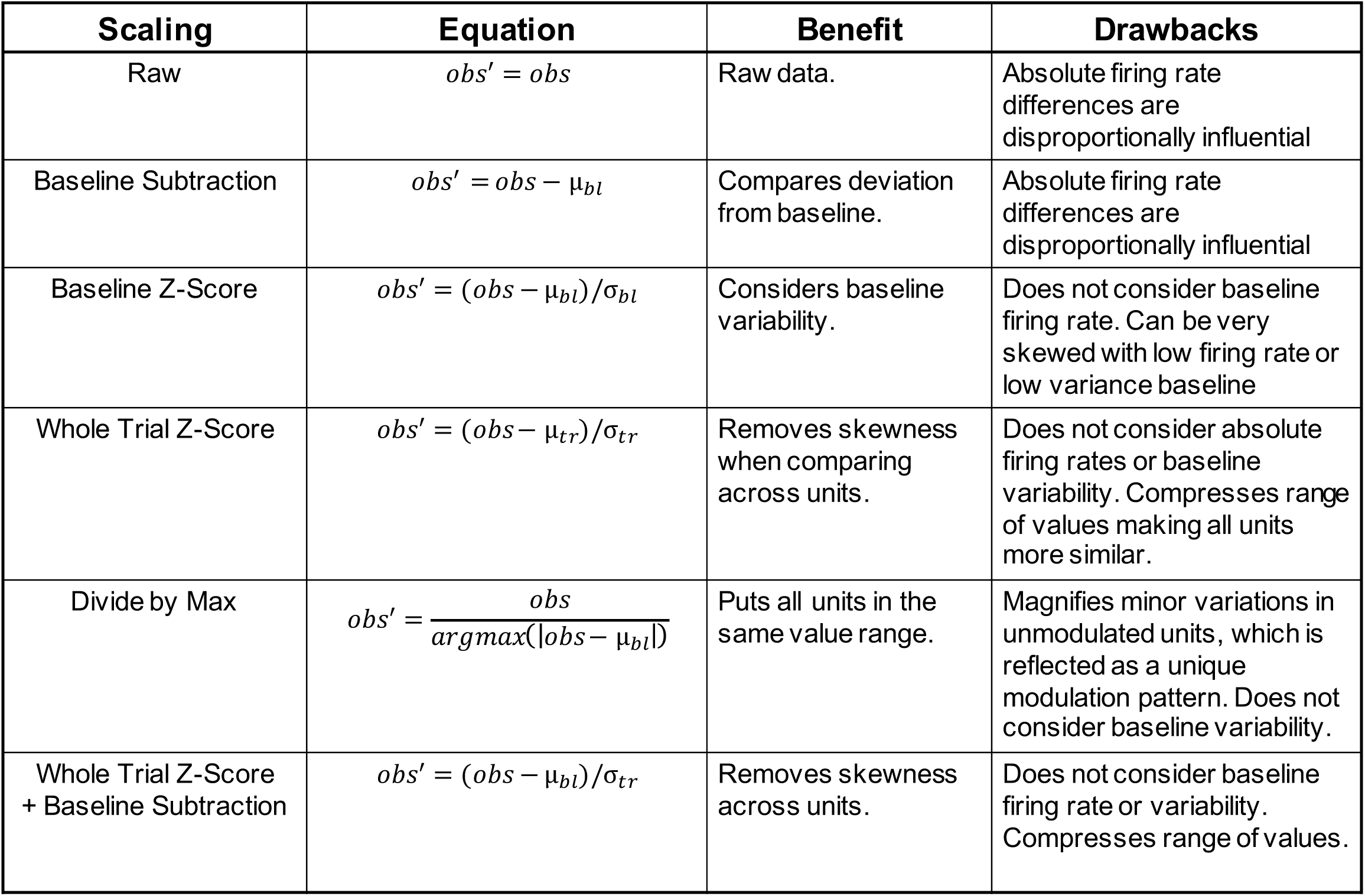
**Scaling procedures.** Six scaling procedures were used across the analyses presented. Each scaling procedure has a theoretical motivation, but at a cost. These costs and benefits are described along with the equation of the scaling and the short-hand name.

**Figure 1:**
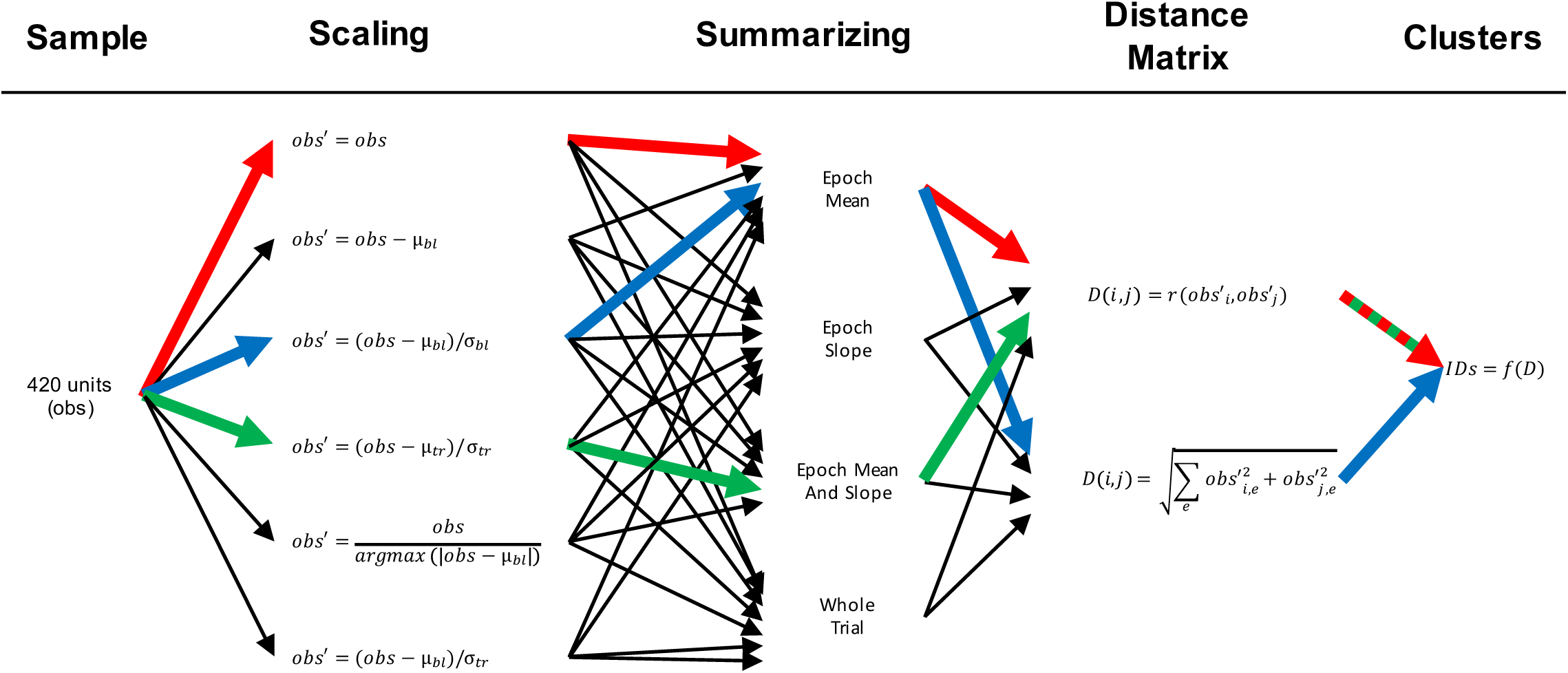
Analysis measures for representative neuron recorded during memory-guided saccade task. Several sets of clustering parameters were used to create a distance matrix used in the classification of observations. The six scaling procedures, four summarization procedures, and two distance measurements are shown. All combinations of parameters were tested, as indicated by the arrows, but in every case the same algorithm *f* was applied to the distance matrix to obtain category identifications. The sets of parameters indicated in blue, red, and green were used for cluster analyses 1, 2, and 3 respectively, and the results are depicted in figures 4, 5, and 6.

This procedure was repeated until all observations were agglomerated into a single group. Then, category identifications were made for a range of number of categories, *k,* by finding the most recent step in the algorithm at which *k* categories with at least *x* members were present. For example, for a *k* of 5, the most recent set with 5 categories of at least *x* members was assigned as the final classification. Category membership for *k* was assessed between 1 and 20. The value of *x* was set to 10; only categories with 10 or more members were considered to assure robust results.

Category membership was assessed for six scaling procedures, four methods of summarizing the response, and two similarity metrics (Table 1; Fig. 1). The merits and drawbacks of the scaling procedures are summarized in Table 1. The mean skewness across time points was used to assess the quality of scaling for cross-unit comparisons. We refer to each combination of scaling procedure, SDF summarization, and similarity metric as a pre-processing pipeline. We evaluated clusters derived from each pipeline but will show outcomes for only three.

Modulation of discharge rates was summarized according to the as following approaches. Three (epoch mean, epoch slope, epoch mean and epoch slope) account for the firing rates during epochs: −200 to −100 ms before stimulus onset, 50 to 100 ms post-stimulus, 100 to 150 ms post-stimulus, −100 to −50ms before saccade, −50 to 0ms before saccade, and 50-100 ms post-saccade. “Epoch mean” summarization was based on the mean firing rates in these epochs, “epoch slopes” summarization was based on the slope of the firing rate changes in these epochs, and “epoch mean and epoch slopes” were based on the concatenation of mean firing rate and slope. “Whole trial” summarization did not parse the responses into epochs, and was instead based on the values of the SDF aligned on stimulus onset (-200 to 300 ms) or saccade (-300 to 200 ms) at each time point to millisecond resolution. To emphasize equally all epochs, means, and slopes, each summary value was individually converted to a Z-score across the sample.

Pairwise distance was measured two ways: Euclidean distance and correlation. Euclidean distance can be conceptualized as the physical distance between two points in multidimensional space while disregarding which dimensions contribute to this distance. Correlation assesses the relationships between the dimensions while disregarding the particular values of those dimensions. The different emphases of these two distance metrics can and often does assign two units to the same category via one metric but not the other (Fig. 2).

**Figure 2:**
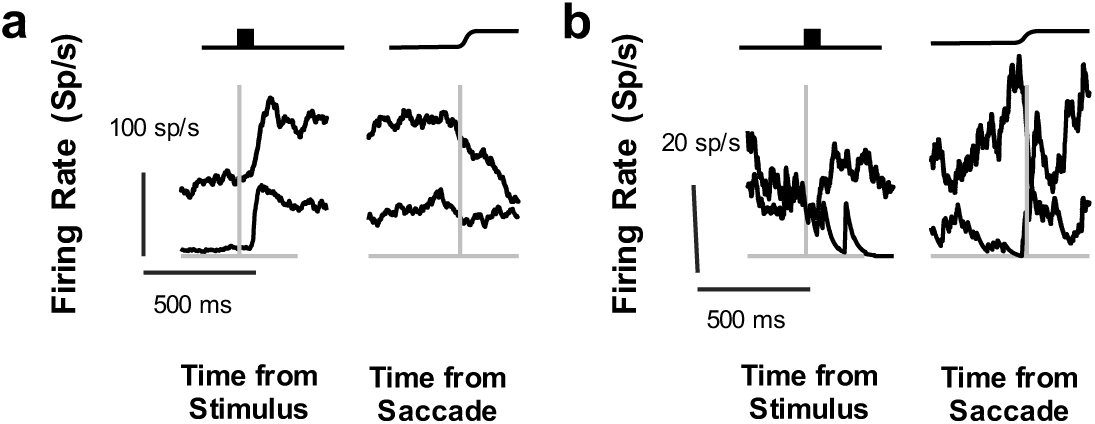
Distance measurement comparison. Two distance measurements were used in the present set of analyses to capture different aspects of cross-unit comparisons. (a) Two example units whose modulation patterns are similar but have different absolute firing rates. These units would have a very small correlation distance and a large Euclidean distance. The left panel shows the average spike density function aligned on stimulus onset, as indicated by the graphic above. The right panel shows the average spike density function aligned on saccade, as indicated by the graphic above. Scale bars are shown in the lower left. (b) Two example units whose modulation patterns are different but have similar absolute firing rates. These units would have a large correlation distance and a small Euclidean distance. Conventions as in (a).

**Euclidean distance.** Based on the firing rate each unit was placed in a multi-dimensional space. This multi-dimensional space had either 6 (for epoch mean and epoch slope summarizations), 12 (for epoch mean and slope summarization), or 1002 (for whole-trial summarization). The Euclidean distance between the units in this space:

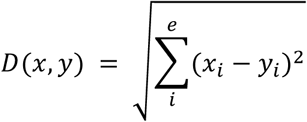

defines a pairwise distance matrix, where *e* is the number of epochs (or milliseconds).

**Correlation.** Based on the firing rate each unit was defined a 6-, 12-, or 1002-element vector, depending on summarization method, and the correlation between two vectors:

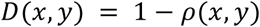

measures the similarity of the modulation patterns of two units while disregarding absolute differences in firing rates.

All of these pre-processing pipelines were tested, and all produced unique results. Some pipelines produced subjectively good clustering, others produced subjectively poor clustering. The results of three representative pipelines are presented, two that produced poor categorizations, for different reasons, and one that produced good categorizations. These three pipelines are presented by color in Figure 1. The first combination uses baseline z-score scaling, summarizes responses based on epoch means, and uses Euclidean distance; the second does not scale the data before summarization, summarizes responses based on epoch means, summarizes responses based on epoch means, and uses correlation distance; the third uses whole-trial z-score scaling, and summarizes responses based on epoch means and epoch slopes.

### Consensus clustering

No *a priori* reason endorses the categories yielded by a particular SDF scaling procedure, summary of response modulation, or distance measurement. Therefore, we employed a novel strategy of combining the results of multiple clustering pipelines. This algorithm considers the pairwise distance between units for each individual pre-processing pipeline tested by creating a composite distance matrix. Each individual distance matrix was z-scored internally to correct for the absolute scale differences between different scaling procedures and distance measurements. Then, the median for each pair was selected to prevent skewing by non-optimal pipelines. Thus, if the pairwise distance between two units was consistently small, then this composite measure was also small, whereas if the pairwise distance between two units was small in some parameter sets but was generally larger, then the composite distance metric reflected the trend toward differences but accounted for the isolated cases of similarity. After creating this composite distance matrix, the same agglomerative algorithm used for each individual pipeline was applied to identify categories. Intuitively, this method was intended to distinguish units that were clustered together regardless of pre-processing pipeline from units that were members of different clusters regardless of pre-processing pipeline.

In essence, this procedure performs a clustering that operates on a distance matrix whose entries represent robustness of categorization across a number of individual procedures, or a “consensus clustering” problem (Goder and Filkov, 2008). Indeed, consensus clustering has been used to identify biophysical classes of neurons (Ardid et al., 2015). However, while conceptually similar, the previous “meta-clustering” and the present consensus clustering differ operationally in both the algorithm used for performing clustering and the pre-processing of input data. Ardid and colleagues performed K-means clustering, which does not provide a unique clustering solution and is highly sensitive to starting points. (Bradley and Fayyad, 1998; Peña et al., 1999; Celebi and Kingravi, 2012), so their meta-clustering involved multiple iterations of the K-means procedure using the same input data and then assigning clusters via robust co-membership across each iteration. Unlike Ardid and colleagues, we used agglomerative clustering, which delivers unique solutions because no optimization steps are involved. However, we found that clustering outcomes were sensitive to the pre-processing pipeline, varying with discharge rate scaling procedure, SDF summarization, and distance measurement. Hence, our consensus clustering approach was conceived to assess cluster assignment consistency over pre-processing steps, not local solutions to clustering one set of pre-processed data.

### Assessing Number of Categories

The number of categories in individual clustering procedures was selected using a lenient version of Tibshirani’s gap procedure (Tibshirani et al., 2001), which assesses the reduction in intra-cluster distance with respect to randomized null-sets created with no intrinsic clustering. Valid splitting of clusters should have a greater than chance reduction in intra-cluster distance, assessed by the standard deviations of the intra-cluster distance in the null sets, whereas excessive splitting should have a reduction in intra-cluster distance within the standard deviation of the null set. The strict version selects *k* categories as the first number of categories meeting this criterion. In the lenient version of this test, each occasion on which the above criterion is met was treated as potentially valid, and visual inspection was used to determine whether categorizations were either insufficient or excessive. In some cases, due to the difficulty in creating a reasonable null-set from physiological data, categories were selected based on the properties of the gap curve. When a reasonable null-set could not be determined, an inflection in the gap-curve was identified. This inflection identified the number of categories at which the reduction of intra-cluster distance was markedly less than that of the previous sequence of clusters.

For consensus clusters and the composite distance matrix, the means of creating a null set in order to use Tibshirani’s gap procedure is unclear. Therefore, in such cases a pair of criteria for determining the maximum number of clusters was set: no more than 10% of units were allowed to remain un-clustered, and each cluster required at least 10 members. The maximum number of categories that met both criteria was selected.

### Comparing Classification Schemes

The quality of alternative categorization schemes was assessed by calculating an index of member variability through the ratio of variances of the spike density values. Specifically, for each time point in the spike density functions, the within-cluster variance at that time point was divided by the variance of the cluster mean across time points. For each cluster, the average ratio was calculated, and then the grand average was taken. Because this ratio will decrease by definition as more clusters are formed, a penalty for over-splitting was imposed by multiplying the grand average ratio by the square root of the number of clusters. That is, for a given cluster the modulation strength was calculated as:

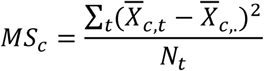

where *c* indexes cluster and *t*, time. Then, for each time point a ratio of variances (*RoV*) was calculated:

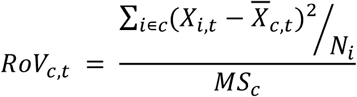

These are then combined by averaging the values for each cluster over time, then across clusters, then applying a penalty for over-clustering proportional to the square root of the number of clusters identified:

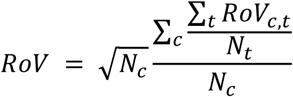

Small *RoV* values can be obtained either through large group-wise modulations over time or a lack of variability among the categories’ constituent members, whereas large *RoV* values are obtained through weak category-wise modulations or large variability. That is, smaller *RoV* values indicate better categorization, and vice versa. No benchmarks have been established for this index, so we interpret relative values in comparing the quality of two categorization schemes.

The similarity of two categorization schemes was assessed by the Adjusted Rand Index (*ARI*) (Hubert & Arabie, 1985):

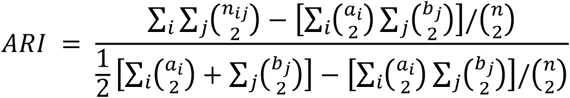

Where *a*_*i*_ and *b*_*j*_ are the counts of category *i* or *j* in categorization procedure *a* or *b*, respectively, and *n*_*i,j*_ is the number of observations in both category *i* in categorization scheme *a* and in category *j* in categorization scheme *b*. This quantity measures similarity of two data categorizations and is adjusted by chance co-categorization produced through the two schemes. To assess significance, each categorization was randomly shuffled separately, destroying internal structure between the two schemes, and *ARI* was recalculated. This was repeated 1000 times and *p* was the proportion of shuffled *ARI* that exceeded the non-shuffled *ARI*.

To visualize the overlap, for each pairwise categorization combination a *signed χ*^*2*^ was calculated:

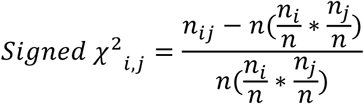

That is, the difference between the observed pairwise count and the count expected by the marginal probabilities of each individual categorization scheme was calculated, then normalized by that expected value. Each category assignment was shuffled separately 1000 times and the signed χ^2^ was recomputed for each combination and iteration. A bootstrapped z-score was then calculated for each category combination.

## RESULTS

This analysis is based on 466 units sampled in FEF from three macaque monkeys performing visually guided and memory guided saccades in pursuit of other research aims.

### Traditional Response Categorization

First, units were categorized based on traditional criteria (Bruce and Goldberg, 1985; Schall, 1991) and the canonical visual, movement, and visuomovement units were identified (Fig. 3). While the average discharge rates of these categories were as expected, the SDF of the individual units categorized into each type exhibited considerable variation. For reference with subsequent analyses, the *RoV* value was 50.89. Thus, while the traditional categorization methods captured general trends in the modulation patterns of FEF neurons, additional variation was present but unaccounted for.

**Figure 3:**
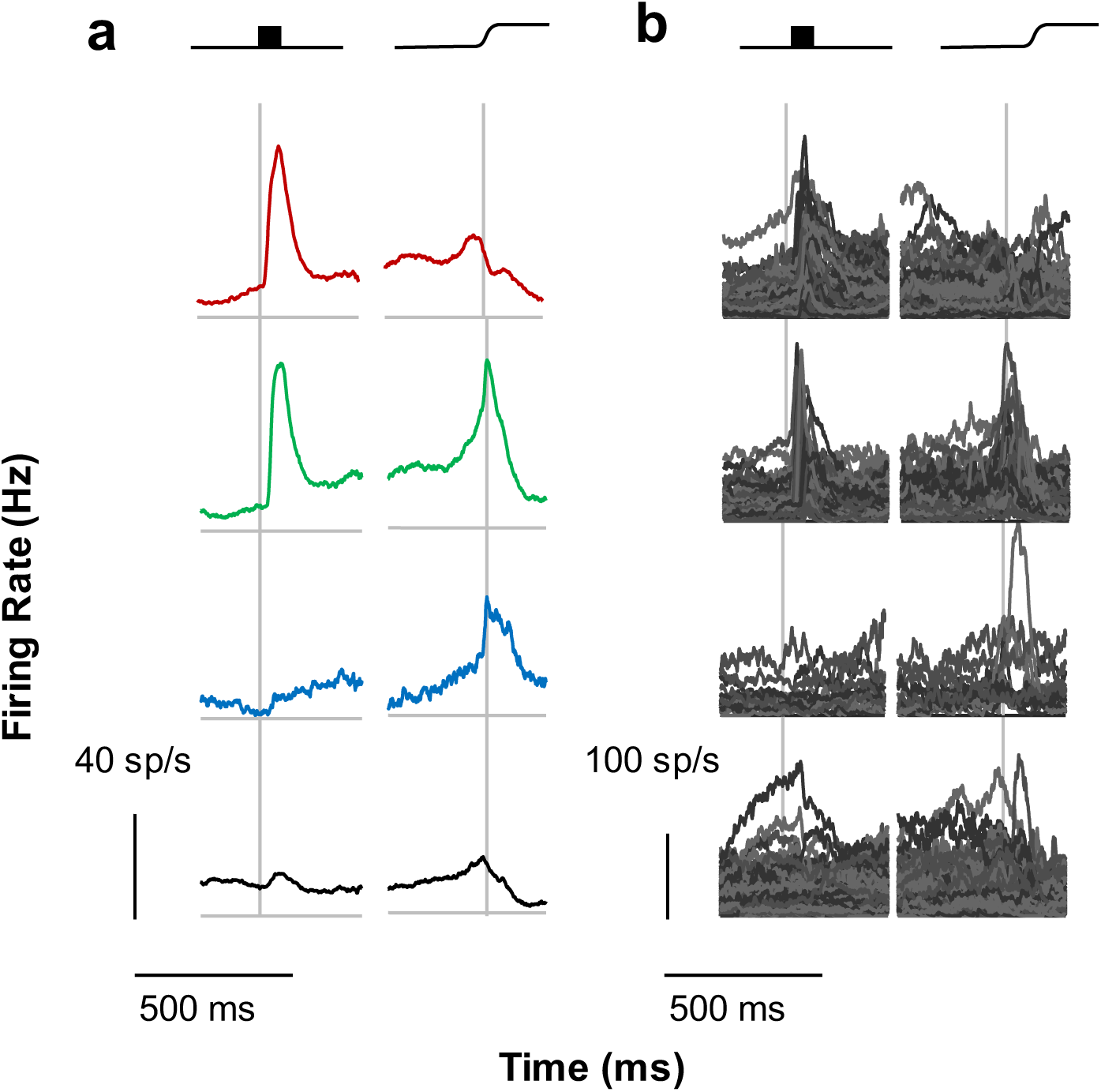
Traditional Classification. The current sample was classified according to traditional criteria. (a) Group mean SDFs for visual, visuomovement, movement, and unclassified neurons are depicted from top to bottom, with left panels aligned on stimulus onset and right panels aligned on saccade. Here and in subsequent figures the categories of neurons are arranged on a visual to motor axis, and colors assigned such that red indicates visual activity and no movement activity, green indicates both visual and movement activity, and blue indicates movement activity without visual activity. Black indicates unclassified neurons. (b) Individual spike density functions comprising each category.

### Cluster Analysis 1: Baseline Z-Score, Epoch Means, Euclidean Distance

To begin accounting for this excessive variability, a cluster analysis was performed on the SDF that were Z-scored based on the mean and standard deviation of the baseline firing rate. This captured the definition of visual and movement activity used above. However, the data-driven clustering procedure revealed functional categories that are similar in their firing rate modulations in more than just two epochs. Based on the mean firing rate in the six specific task epochs described in Methods, each unit was represented as a six-element vector. Based on Euclidean distance measures of pairwise distances 8 categories of units, numbered 1_a_-8_a_ to distinguish this set of results, were found (Fig. 4). Unlike the traditional categorization scheme, two categories of visual units were identified, identified as categories 1_a_ and 4_a_. Both categories had modest visual responses and no perisaccadic activity. They were differentiated by the presence or absence of anticipatory activity before the target appeared and by delay period activity. An additional category, category 5_a_, had a robust visual response with weak presaccadic ramping. Four categories of units had both visual and presaccadic responses. Two of these categories had robust visual responses and intermediate presaccadic ramping: categories 2_a_ and 7_a_. These categories were differentiated by the return to baseline after the saccade: category 7_a_ had a typical slow return to baseline whereas category 2_a_ returned to baseline almost immediately. The other two categories, 6_a_ and 8_a_, had only modest firing rate modulation in both epochs and were distinguished by the presence (8_a_) or absence (6_a_) of anticipatory activity. The final category, 3_a_, did not show firing rate modulation during the trial. This demonstrates that additional diversity is present in FEF firing patterns that has been unaccounted for in the traditional scheme. However, this clustering approach failed to identify purely movement neurons or post-saccadic neurons.

**Figure 4:**
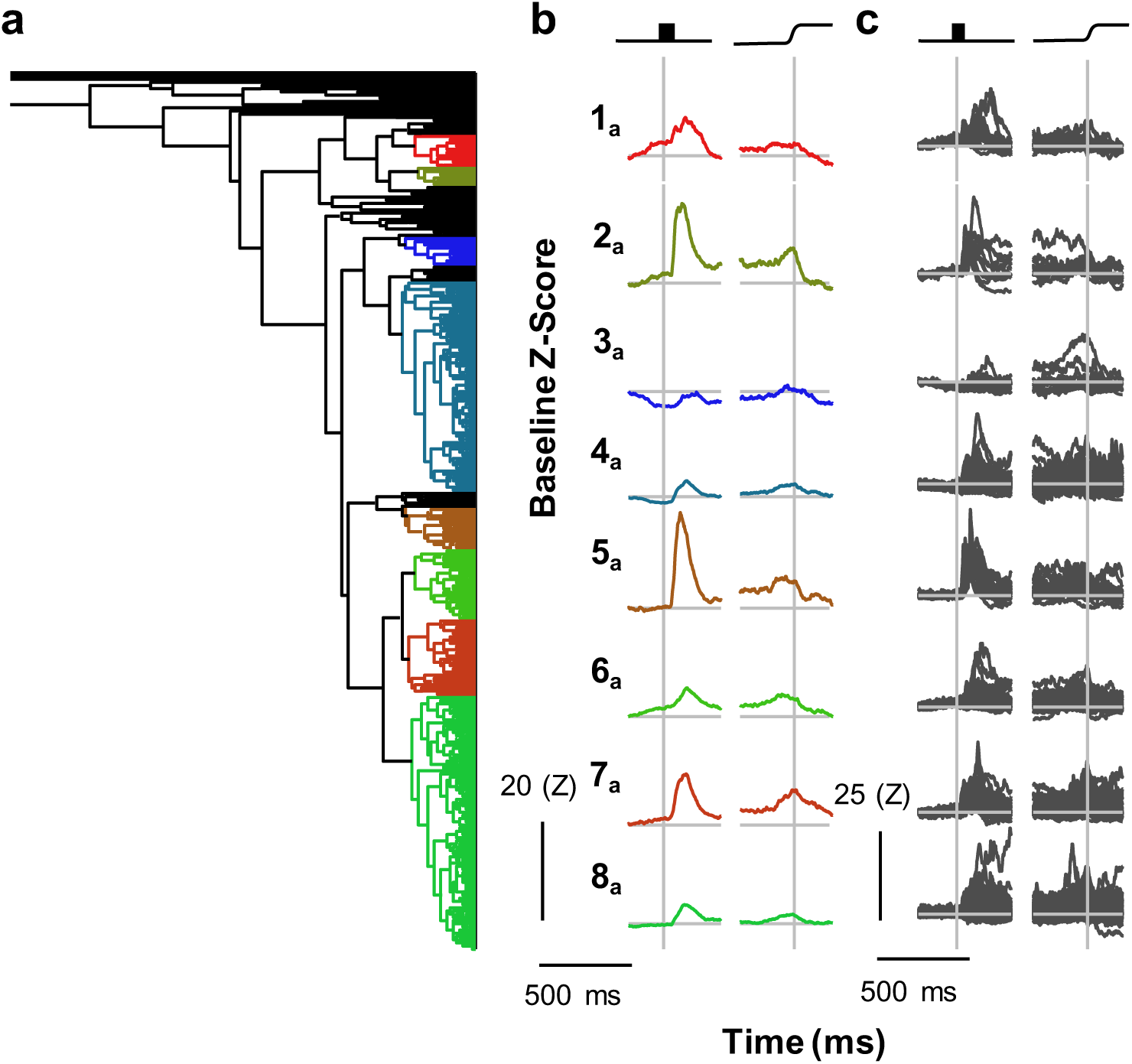
Cluster Analysis 1. Neurons were categorized via cluster analysis using the baseline z-score scaling procedure, epoch mean summarization, and Euclidean distance. (a) The dendrogram resulting from cluster analysis 1 shows the eight identified categories. Horizontal distance indicates pairwise similarity with individual neurons on the right and full agglomeration on the left. Colors indicate categories and are arbitrarily assigned on a visual to motor axis as in Figure 3. (b) Category means are plotted aligned on stimulus onset and saccade. Each category was given an arbitrary numerical identifier for convenience. Scale bars are shown in the lower left. (c) Individual neurons comprising each category aligned on stimulus and saccade. Scale bars are shown in the lower left.

The eight categories did not match the three traditional categories in FEF that should have been recovered through cluster analysis. However, similar to the traditional categorization scheme, these category means captured some general trends of the individual units comprising the categories. Variation was reduced (*RoV* = 18.36 relative to 50.89 for the traditional classification), but considerable variation was still evident. This was particularly pronounced in categories 4_a_ and 8_a_ which were also much larger categories than the other six. Thus, these categories seem to be “catch-all” categories. Other categories seem to be nearly identical, though are clearly seen as different groups in the dendrogram (e.g., 2_a_ and 5_a_). The dendrogram in Fig. 4a also shows that units did not exhibit clear clustering. Instead, it appears as though small groups or individual units are progressively grouped together such that, at the clustering step when eight categories meet the membership criterion, 16% of units were still uncategorized. That is, for the number of clusters identified via Tibshirani’s gap procedure, 16% of neurons were so dissimilar to each other and the eight categories that they could not be placed in any of the eight categories, or form a ninth separately. This may be so because the variation of the SDF of the identified units was high (mean skewness = 1.35). Taken together, these considerations indicate that this clustering procedure is insufficient.

### Cluster Analysis 2: Non-normalized SDF, Epoch Means, Correlation Distance

To account for more of the excessive variability, a cluster analysis using a correlation distance measurement was performed on the non-normalized data. This approach captures relative rather than absolute changes in firing rate. That is, if two units were similarly modulated but have different firing rates, this procedure treated them as members of the same category. In other words, this approach emphasized the pattern of modulation of FEF neurons rather than the absolute discharge rate.

This procedure identified 6 categories, 1_b_-6_b_ (Fig. 5). These categories did not match the three traditional categories; movement neurons are missing. Instead, each of the six identified categories demonstrated modulation following visual stimulation to different degrees. Two of these categories, 1_b_ and 6_b_, showed visual modulation only, and were differentiated by the baseline firing rate and degree of visual modulation. Three categories had both visual and movement-related activity: 2_b_, 4_b_, and 5_b_. Category 2_b_ had modest visual modulation and no delay activity; category 4_b_ had modest visual activity and some delay activity, and category 5_b_ had robust visual modulation, prominent delay activity and presaccadic ramping, but its activity fell off dramatically at the time of the saccade. The final category, 3_b_, had modest visual modulation and a sharp post-saccadic transient.

**Figure 5:**
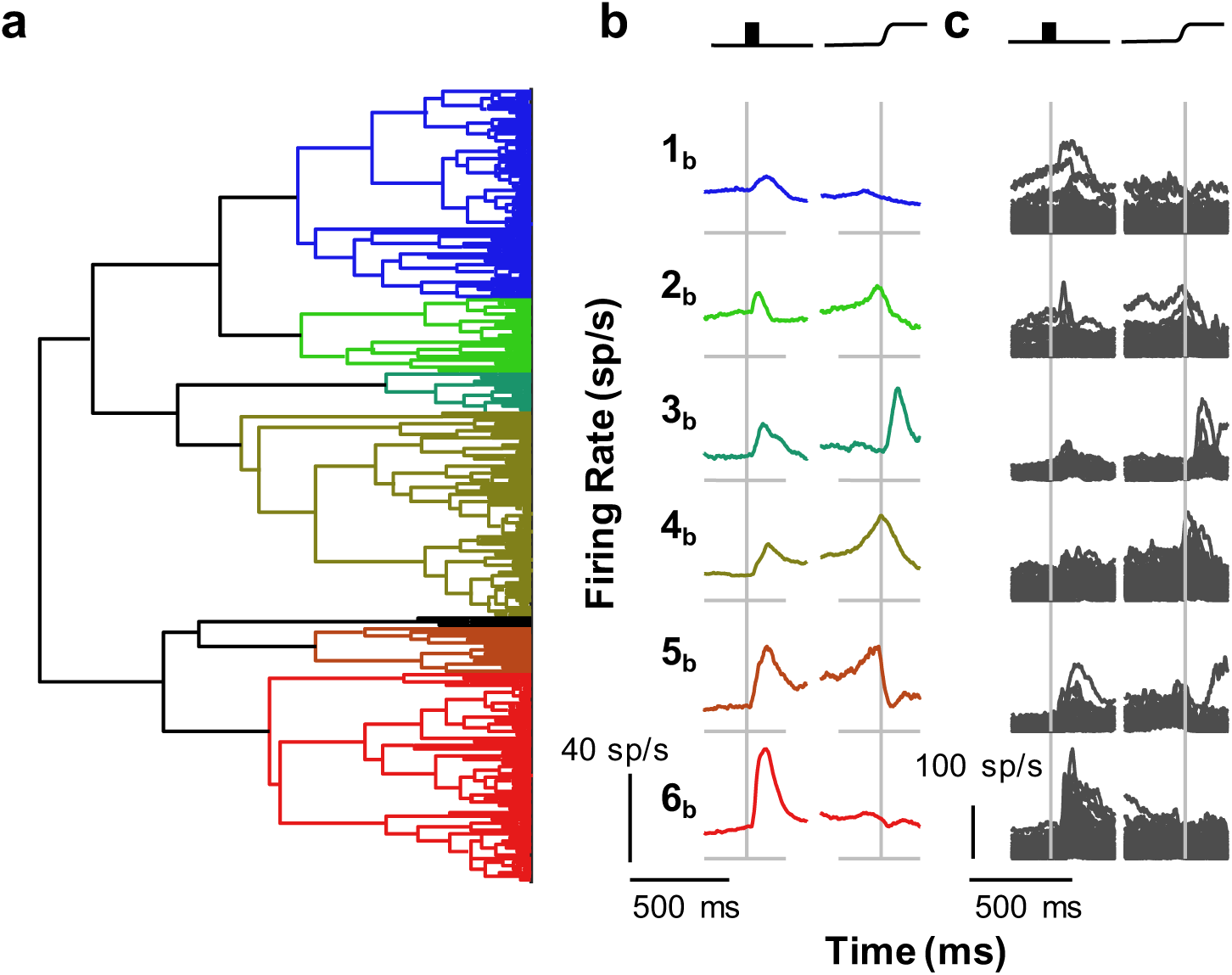
Cluster Analysis 2. Neurons were categorized via cluster analysis using no scaling procedure, epoch mean summarization, and correlation distance. Conventions for (a) through (c) as in Figure 4.

Considerable variability within categories remained and in fact increased relative to cluster analysis 1 (*RoV* = 29.70 as opposed to 18.36), though the category means did capture category trends. This variability may be due to the skewed firing rates that range from 0 to 210 sp/s (mean skewness: 2.09) which allowed few units in one category to drive modulations apparent in the category means. For example, the inclusion of some units with large visual responses and high firing rates in category 2_b_ was a driving factor in the modest visual activity seen in the category mean. Thus, both the baseline z-score and un-normalized firing rates may be ineffective for comparing across units, even though they are useful for assessing within-unit modulation. However, the appearance of the dendrogram for this clustering outcome should be noted (Fig. 5a). Unlike that from analysis 1, a more sensible structure is apparent when this combination of clustering parameters was used.

### Cluster Analysis 3: Whole Trial Z-Score, Epoch Means and Slopes, Correlation Distance

To account for more of the variability in modulation patterns, a different scaling procedure was used: Z-scoring across the entire trial and summarizing the SDF with both the means of the SDF and the slopes during the relevant epochs were considered. The agglomerative clustering algorithm identified five categories, 1_c_-5_c_ (Fig. 6). Three of these categories had visual activity only: 1_c_, 2_c_, and 4_c_. Category 4_c_ had pronounced delay activity. The other two were distinguished by the time of peak visual response, with the visual activity of category 1_c_ peaking earlier and that of category 2_c_, later. Category 3_c_ had robust visual and presaccadic modulation but did not have delay activity. The final category, 5_c_, showed robust presaccadic ramping and only modest, if any, visual activity. It should be noted that the presaccadic ramping activity in category 5_c_ peaked after the saccade and showed a slow reduction of firing rate back to baseline, whereas the category with both visual and saccadic responses had a peak perisaccadic activity at the time of the saccade followed by a sharp return to baseline, but this sharp return is not as pronounced as the “clipped” movement neurons in category 5_b_. This indicates that the additional diversity evident in visual inspection of discharge rate modulation patterns is tangible and identifiable. Further, of the 3 cluster analyses, analysis 3 produced the classification most similar to the traditional. Visual and visuomovement categories were identified as well as a putative pure movement group (category 5_c_).

**Figure 6:**
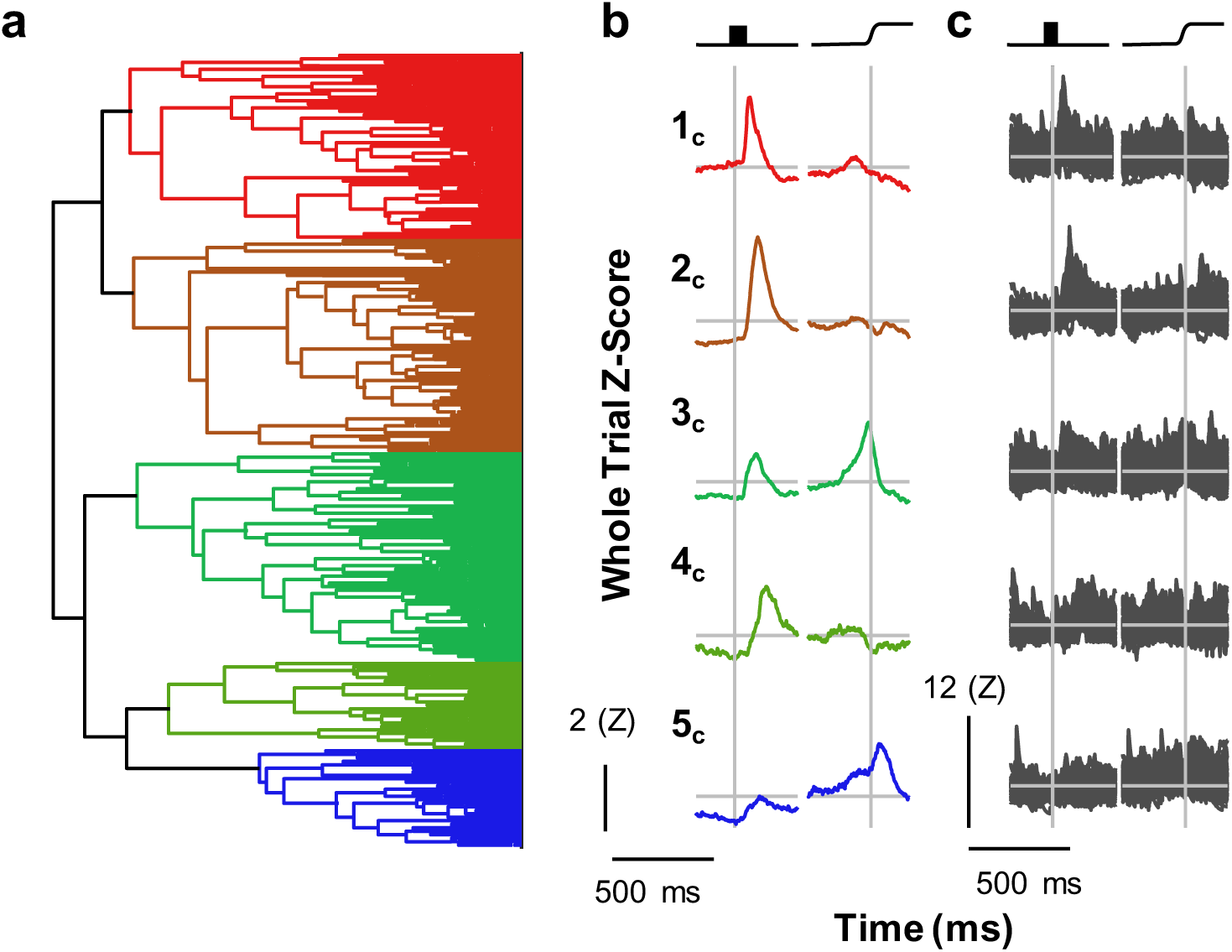
Cluster Analysis 3. Neurons were categorized via cluster analysis using the whole trial z-score scaling procedure, epoch mean and slope summarization, and correlation distance. Conventions for (a) through (c) as in Figure 4.

The range of values through this normalization was smaller and is less skewed (range: −3.3–7.6; mean skewness: 0.73), suggesting that this scaling method provided a more equitable cross-unit comparison. As with analysis 2, these categories are apparent in the dendrogram structure, though some additional heterogeneity can be observed, particularly among categories 1_c_ and 2_c_. This may explain, in their particular cases, the seeming similarity between the two; it could be that splitting further would reveal additional heterogeneity, and in the case of the category 3_c_, further splitting could reveal a second visuomovement group without “clipped” activity as well as a pure movement category. However it should be noted that unlike cluster analyses 1 and 2, the category means were much more representative of the members (*RoV* = 5.19 as opposed to 18.36 and 29.70 for cluster analyses 1 and 2, respectively).

### Consensus clustering

We now address the problem of individual units being members of different categories following different analysis paths (Fig. 7). This occurs because different pre-processing pipelines and distance metrics resulted in different distance matrices upon which the agglomerative clustering algorithm operates. Consequently, a given pair of units were members of the same category following one pipeline but members of different categories following another pipeline. However, no particular pipeline was more confidently motivated or more certainly correct than another. Nevertheless, assuming the existence of ground-truth categories, consistent with anatomical constraints, units that are actually members of the same ground-truth category should have small pair-wise distances regardless of scaling or clustering procedure. Likewise, units that are members of different ground-truth categories would have small pair-wise distances only as an artifact of particular measurement parameters and clustering algorithm.

**Figure 7:**
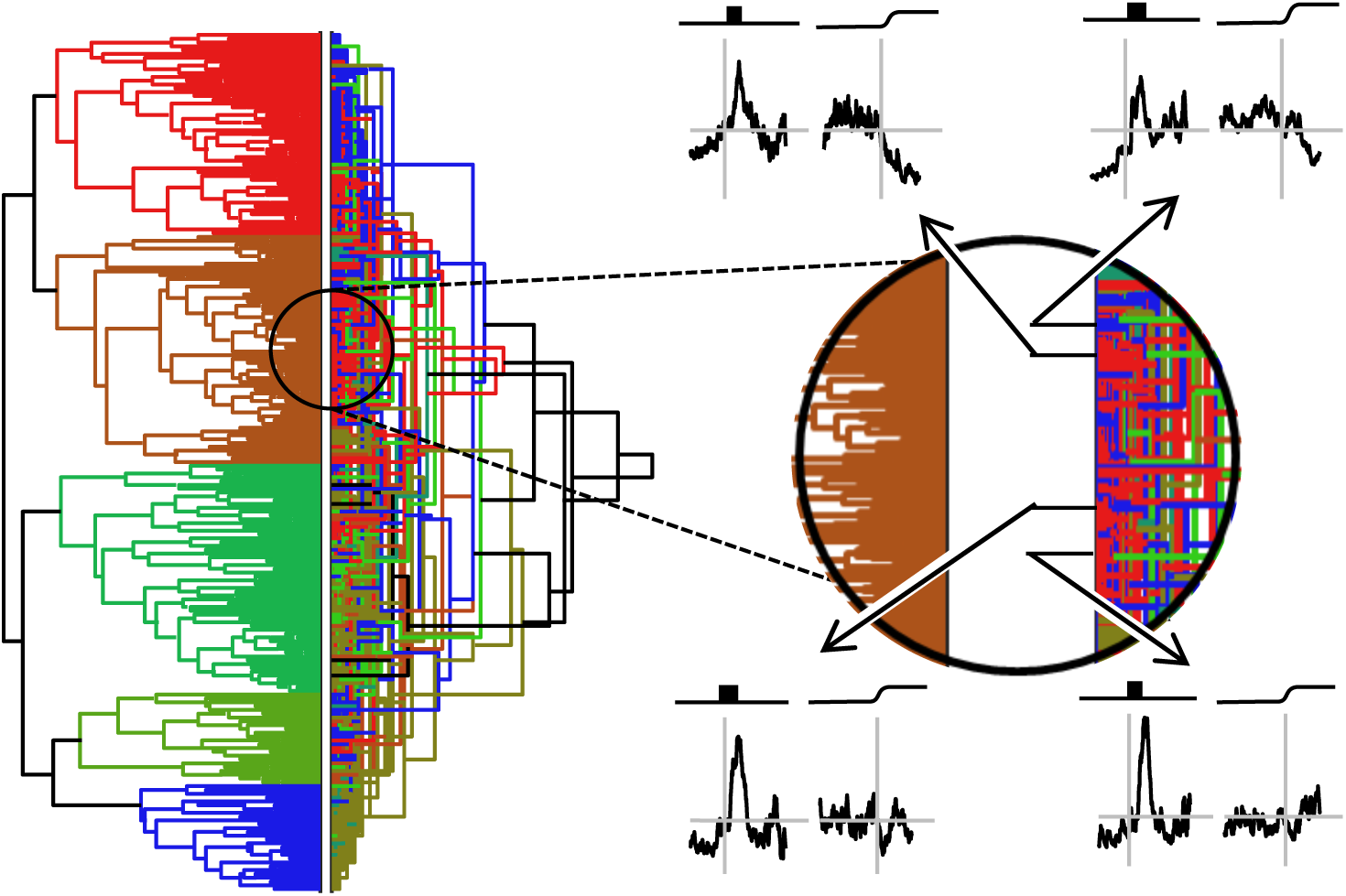
Comparison of Cluster Analyses 2 and 3. Left: The dendrograms from cluster analysis 2 and 3 are shown side by side. The dendrogram for cluster analysis 2, right, was vertically re-ordered such that the abutting neurons are the same. In general, the visually related categories indicated by reds and oranges align in both dendrograms, as do the movement related categories indicated by blues and greens. Right: A closer examination of four units. All four units were placed in category 2c. Three were placed in category 6b whereas one was placed in category 1b. Thus the two analyses provide similar, but not identical, categorizations, but there is no principled way of deciding which is correct.

To address this fundamental problem, we employed a second-order clustering procedure known as “consensus clustering” (Strehl and Ghosh, 2002; Goder and Filkov, 2008). We created a composite distance matrix by z-scoring individual distance matrices from all of the pre-processing pipelines (Fig. 1), then calculating the median distance across all parameter sets. This composite distance metric was used to identify units that were consistently similar to one another across pre-processing pipelines and clustering algorithms. The agglomerative clustering algorithm was applied to this distance matrix to identify robust categories of units superordinate to any individual cluster analysis. The last splitting producing a cluster meeting membership criteria was selected for determining the final number of categories.

Consensus clustering identified 10 categories, clearly distinguished in the dendrogram and evident in the distance matrix (Fig. 8). These categories were robust and consistent (*RoV*: 3.91). Even with the penalty for over-clustering in the *RoV* metric, consensus clusters account for more of the variability in the neural data than the classification produced by the best individual classification.

**Figure 8:**
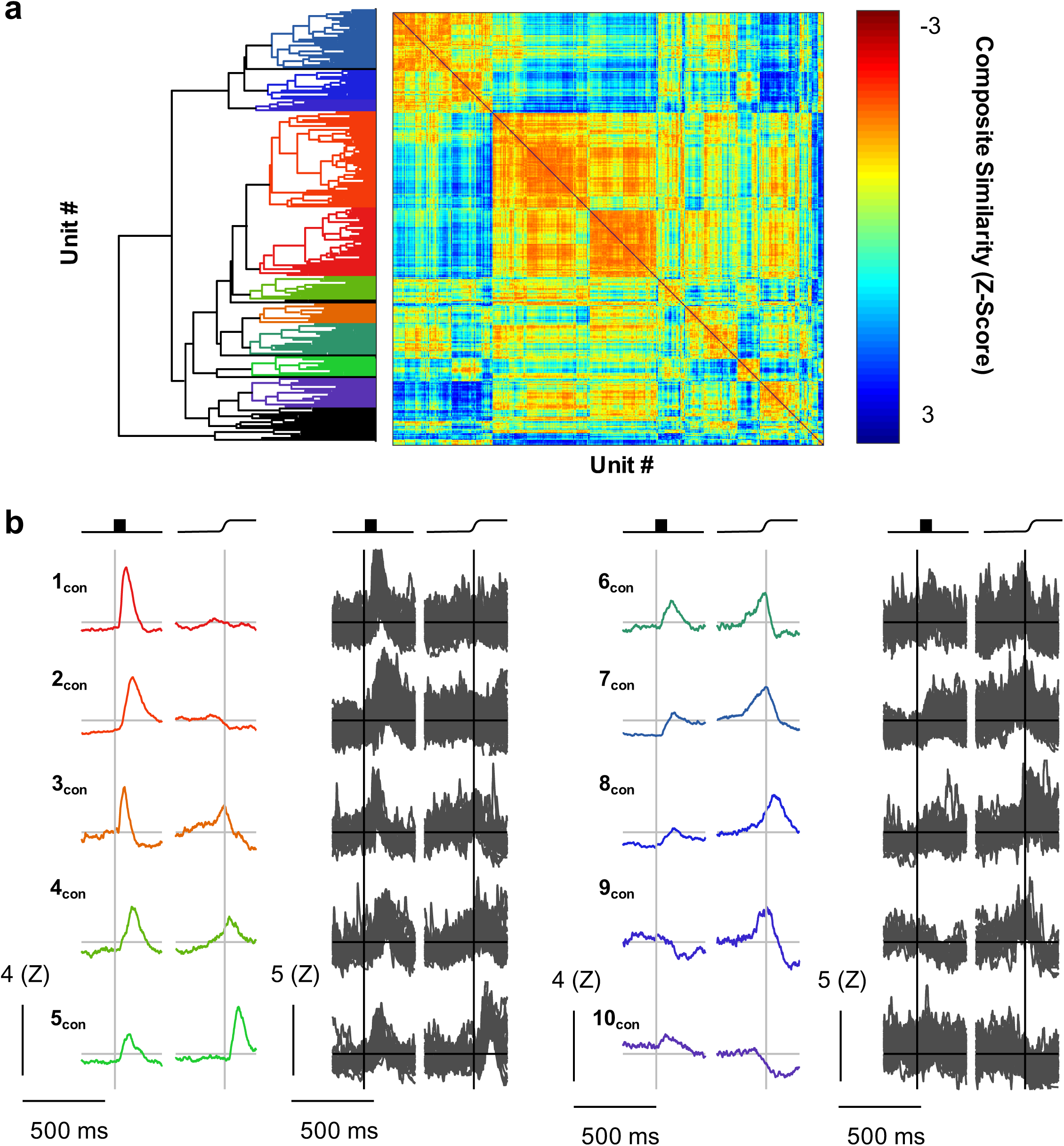
Consensus clusters. Consensus clusters were identified by creating a composite distance matrix and applying the agglomerative clustering procedure to this matrix. (a) The resulting dendrogram is shown abutting the composite distance matrix. Colors in the dendrogram are assigned as in Figs. 3-6. Color in the distance matrix is indicative of composite similarity. Warm colors (low composite z-score) indicate consistently similar units whereas cool colors indicate consistently different units (high composite z-score). (b) The category mean SDFs (columns 1 and 3) and the individual SDFs (columns 2 and 4) comprising them are shown aligned on stimulus onset (right) and saccade (left). Scale bars are shown in the lower left of each column. Arbitrary category labels were assigned for convenience.

#### Categories with visual responses only

We identified two categories of neurons that had visual, but not saccade-related, activity. These categories, 1_con_ and 2_con_, showed flat baseline activity and a sharp visual transient. The time of peak firing rate differentiated these two categories, with mean peak latencies of 74 ms (1_con_) and 136 ms (2_con_). Also, category 2_con_ had persistent delay activity until the saccade.

#### Categories with visual and saccade-related facilitation

We identified five categories of neurons with both visual and pre-saccadic increases in firing rate. Two of these categories, 3_con_ and 4_con_, showed marked increases in firing rates following visual stimulation and were distinguished by the time of peak visual activity (mean values of 70 ms and 161 ms respectively). They were also distinguished by the time and character of the pre-saccadic ramping. The firing rate of the category 3_con_ peaked at the time of the saccade and quickly returned to baseline whereas the firing rate of category 4_con_ peaked after the saccade and returned to baseline more slowly. Two of the three remaining categories, 6_con_ and 7_con_, also had clipped movement activity with late, weak visual responses. These two categories are differentiated by the absence (6_con_) or presence (7_con_) of delay period activity. The final category with both visual and movement activity, 8_con_, showed only modest visual activity and may be more accurately described as a pure movement category. In either case, the movement activity peaked just after the saccade but returns slowly to baseline. An additional category, 5_con_, was not movement-related *per se*, but exhibited a strong post-saccadic transient with a modest, early visual response.

#### Categories with response attenuation

Two categories of units showed distinct decreases in firing rate. The first of these, 9_con_, was unique in having an “off” response to visual stimulation, but it also showed clipped presaccadic ramping that peaked just before the saccade. This provides a fruitful contrast with the final visuomovement category, 8_con_, which showed only a modest increase in firing rate but robust, un-clipped perisaccadic ramping. This distinction may be a useful criterion in future studies examining the differences in stimulus-driven or goal-directed saccades. The second category with an off response, 10_con_, showed little or no visual modulation, but was sharply inhibited around the time of the saccade, resembling the canonical fixation neurons.

## DISCUSSION

We applied a consensus clustering technique and identified ten robust functional categories in FEF based on modulation of discharge rates alone. This categorization includes but exceeds the traditional categories. First, we will discuss the relationship of the new functional categories to the traditional categorization. Second, we will discuss potential anatomical correlates of these consensus clusters. Finally, we will discuss the limitations and extensions of consensus clustering techniques.

### Correspondence with Traditional Functional Classifications

To compare the traditional and this new categorization, we assessed their overlap by calculating the proportion of consensus clusters identified as visual, movement, visuomovement, or neither. The traditional scheme and our new consensus clustering procedure show significant overlap (Fig. 9; *ARI* = 0.0931, p < 0.001). Thus, rather than contradicting the traditional categorization, this approach offers more fine-grained partitions. In fact, although we did not explicitly test for post-saccadic or fixation neurons, both categories were identified via consensus clustering. This unsupervised discovery increases confidence that consensus clustering identifies natural neural categories.

**Figure 9:**
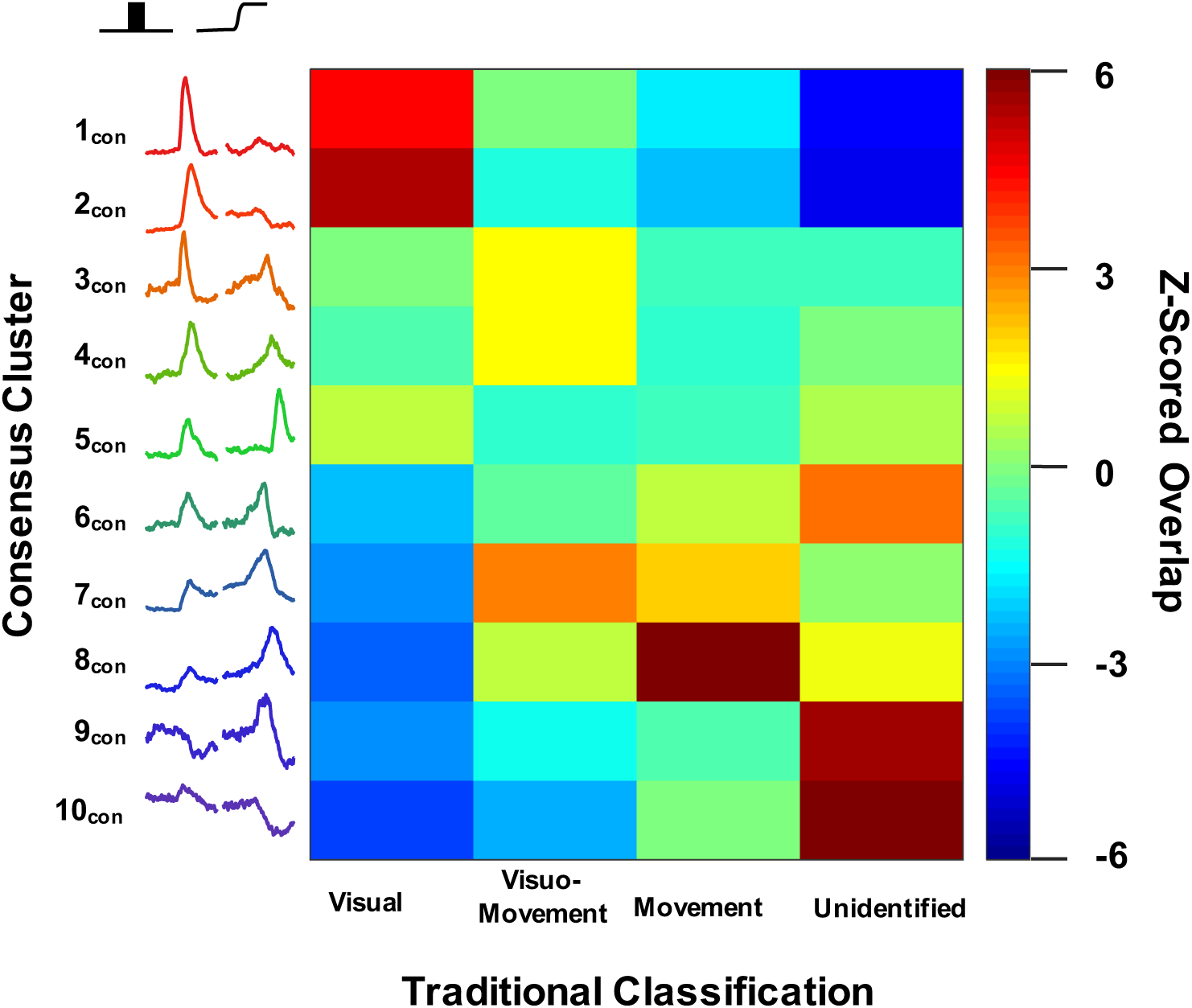
Relation to traditional classification. The consensus cluster assignments were compared to traditional classifications. The consensus clusters are depicted on the vertical axis and traditional classification on the horizontal. The color in the heat map indicates the prevalence of neurons being classified in a given combination. For a given cell in the matrix, a warm color indicates that more neurons were assigned to both that consensus cluster and that traditional classification than expected by chance, green indicates that the expected number of neurons were assigned to both categories, and a cool color indicates that fewer than expected neurons were assigned to both categories. Cluster 1_con_ and 2 _con_ neurons were more often identified as visual cells and were rarely uncategorized. Cluster 3 _con_, 4 _con_, and 7 _con_ neurons were often identified as visuomovement cells. Cluster 8 _con_ neurons were more often identified as movement cells and not visual cells. Cluster 9 _con_ and 10 _con_ neurons were generally not categorized, but when they were they were not classified as visual cells.

The variation captured by this new classification is not surprising. For example, the distinction of transient and phasic visual responses (Cleland et al., 1971) has been evident in FEF (e.g., Sato et al., 2001). Likewise, clipped movement neurons have been associated with saccade dynamics in superior colliculus (Waitzman et al., 1991), but their existence in FEF has been uncertain, being absent in one sample (Segraves and Park, 1993) and present in another (Hanes et al., 1995). Confirming the presence of clipped movement neurons in FEF substantiates the hypothesis that FEF contributes to the dynamics of saccade production (Schiller et al., 1987).

### Correspondence with Anatomical Constraints

Unsupervised clustering can enhance our understanding of the signal processing within FEF and between FEF and other cortical areas and subcortical structures. For example, there is a disagreement regarding the signals conveyed from FEF to SC. One study found only movement and fixation neurons antidromically stimulated from the SC (Segraves and Goldberg, 1987). A subsequent study replicated the movement and fixation signals but found also that visual and visuomovement neurons projected to SC (Sommer and Wurtz, 2000). The disagreement may be resolved by considering more refined categories of neurons.

Another example involves the latency and dynamics of visual responses. Unsupervised clustering distinguished units with early, brisk visual responses from units with later, more sluggish visual responses. Brisk, early responses are characteristic of the M-pathway and are thought to carry spatial location, whereas later and sustained responses are characteristic of the P-pathway and are thought to carry information related to form or object identity (e.g., Nassi and Callaway, 2009). The existence of such categories in FEF indicates that individual units receive segregated input from the different pathways. Further distinctions are likely to be found across the visual eccentricity and saccade amplitude map of FEF given the quantitative and qualitative differences in connectivity of medial and lateral FEF (Barbas and Mesulam, 1981; Barbas and Pandya, 1989; Preuss and Goldman-Rakic, 1991; Schall et al., 1995a; Markov et al., 2014).

### Limitations and Extensions of Clustering Procedures

Each of the individual clustering procedures we used is limited by the distance measure used to calculate pairwise similarity, the method of summarizing the responses, and the quality of the discharge rate samples. Different distance measures emphasize different aspects of the variability across units (Fig. 2). Whereas Euclidean distance emphasizes similarity in absolute discharge rates, correlation distance emphasizes similarity in the pattern of modulation of discharge rates. Different summaries of the variation of discharge rate emphasize different aspects of the variability across units. A summary of the mean firing rate in different epochs captures absolute discharge rates but ignores dynamics. A summary of the slopes of the SDF in different epochs ignores absolute discharge rates. A summary combining means and slopes across epochs or just using the SDF from the entire trial can expose the clustering algorithm to excessive incidental variation. Different approaches to scaling the SDF across units emphasize different aspects of the variability across units. As shown in Figs. 3-5, different methods of scaling the SDF across units can result in category means that do not accurately represent the individuals comprising those categories. Naturally, different scaling procedures emphasize useful information about the units. For example, z-scoring the SDF based on the pre-stimulus baseline activity emphasizes the magnitude of modulation relative to the variation in the baseline. On the other hand, z-scoring the SDF based on the entire trial reduces the skewed variation of discharge rates. Analytical choices must be made, so confidence in the outcome of every particular clustering pipeline can be questioned.

Consensus clustering increases confidence in distinctions identified across measurements and clustering procedures by minimizing spurious classifications arising from incidental analysis choices or unreliable data. Moreover, the consensus clustering approach affords the opportunity to include as many other measures and clustering procedures as desired. In particular, biophysical spiking properties are certainly useful for categorizing neurons. Spike waveform putatively distinguishes anatomical classes, wherein narrow-spiking units are thought to be inhibitory interneurons and broad-spiking units are pyramidal neurons (McCormick et al., 1985; Connors and Gutnick, 1990; Constantinidis and Goldman-Rakic, 2002; Mitchell et al., 2007; Ardid et al., 2015; but see Vigneswaran et al., 2011). We have previously shown that visuomovement neurons have narrower spike widths than do visual or movement neurons (Cohen et al., 2009; see also Ding and Gold, 2012; Thiele et al., 2016). Other measures like mean firing rate, Fano factor, and coefficient of variation distinguish subtypes of broad- and narrow-spiking neurons (Ardid et al., 2015). The eventual inclusion of such features will surely add complexity but should certainly approach an accurate account of the true diversity of functional neuron categories in FEF.

## Acknowledgements

This work was supported by NEI RO1-EY08890, NEI P30-EY008126, U54-HD083211 and by Robin and Richard Patton through the E. Bronson Ingram Chair in Neuroscience. We thank Joshua Cosman and Wolf Zinke for sharing data, Andy Tomarken for statistical advice, Thilo Womelsdorf for discussions of previous versions of the manuscript, and the Vanderbilt Advanced Center for Computing for Research and Education for access to the high-performance computing cluster. Requests for materials should be addressed to JDS.

